# TRPV1 expressed throughout the arterial circulation enables inflammatory vasoconstriction

**DOI:** 10.1101/2020.02.04.934117

**Authors:** Thieu X. Phan, Hoai T. Ton, Hajnalka Gulyás, Róbert Pórszász, Attila Tóth, Rebekah Russo, Matthew W. Kay, Niaz Sahibzada, Gerard P. Ahern

## Abstract

The capsaicin receptor, TRPV1, is a key ion channel involved in inflammatory pain signaling. Although mainly studied in sensory nerves, there are reports of TRPV1 expression in isolated segments of the vasculature, but whether the channel localizes to vascular endothelium or smooth muscle is controversial and the distribution and functional roles of TRPV1 in arteries remain unknown. We mapped functional TRPV1 expression throughout the mouse arterial circulation. Analysis of reporter mouse lines TRPV1^PLAP-nlacZ^ and TRPV1-Cre:tdTomato combined with Ca^2+^ imaging revealed specific localization of TRPV1 to smooth muscle of terminal arterioles in the heart, fat and skeletal muscle. Capsaicin evoked inward currents and raised intracellular Ca^2+^ levels in arterial smooth muscle cells, constricted arterioles *ex vivo* and *in vivo* and increased systemic blood pressure in mice and rats. Further, capsaicin markedly and dose-dependently reduced coronary flow. Pharmacologic and/or genetic disruption of TRPV1 abolished all these effects of capsaicin as well as vasoconstriction triggered by lysophosphatidic acid, a bioactive lipid generated by platelets and atherogenic plaques. Notably, ablation of sensory nerves did not affect the responses to capsaicin revealing a vascular smooth muscle-restricted signaling mechanism. Moreover, unlike in sensory nerves, TRPV1 function in arteries was resistant to activity-induced desensitization. Thus, TRPV1 activation in vascular myocytes of resistance arterioles enables a persistent depolarizing current, leading to constriction of coronary, skeletal muscle, and adipose arterioles and a sustained increase in systemic blood pressure.

## Introduction

Transient Receptor Potential Vanilloid 1 (TRPV1) is perhaps the best-studied member of the TRP ion channel family. The existence of highly specific and potent pharmacological agonists, including capsaicin and resiniferatoxin,^1, 2^ and spider toxins,^3^ has enabled the interrogation of TRPV1 from the molecular to the whole animal level. Furthermore, pioneering high-resolution (∼3.4 Angstroms) electron cryomicroscopy has revealed key structural features of the TRPV1 channel in the un-liganded and bound state.^4-6^ The functional roles of TRPV1 have been appreciated for over 100 years owing to the ability of capsaicin to elicit pronounced pain responses.^1, 2^ TRPV1 is highly expressed in a subset of sensory afferent nerves with cell bodies located in the dorsal root, trigeminal and nodose/jugular ganglia.^7^ In many of these neurons TRPV1 acts as an integrator of noxious thermal and chemical stimuli including elevated heat, protons, and lipid mediators.^2, 8^ These stimuli activate TRPV1, a non-selective cation channel with considerable Ca^2+^ permeability,^7^ to depolarize the membrane and also to trigger the secretion of neuropeptides from nerve endings.^1, 2^ Accordingly, genetic or pharmacologic inhibition of TRPV1 attenuates inflammatory pain.^9-11^ Further, recent studies have revealed additional functions for TRPV1 as a transduction channel in both itch- and warm-temperature coding neurons,^12,13^ thus indicating a broader role in somatosensory transmission.

Outside of sensory nerves the expression and function of TRPV1 remains controversial. While there is considerable evidence for expression in select brain neurons,^14^ whether or not TRPV1 is functionally present in other tissues is less clear. Of note, several studies have described TRPV1 function in arterial smooth muscle cells;^14-16^ confirmed by measuring Ca^2+^ transients or vasoconstriction in response to capsaicin. On the other hand, other studies have located TRPV1 in vascular endothelium,^17, 18^ and indicated that TRPV1 agonists dilate vessels to lower blood pressure. To further complicate matters, TRPV1 in sensory nerves can exert a neurogenic regulation of nearby blood vessels through the release of vasoactive peptides such as calcitonin gene related peptide (CGRP) or substance P. Indeed, local application of capsaicin to the skin is well known to cause a vasodilation response accompanied by edema.^19, 20^

Here we have exploited reporter mice combined with functional analyses to map TRPV1 expression throughout the arterial circulation. We show that TRPV1 is restricted to the smooth muscle of arterioles, notably in skeletal muscle, heart and adipose tissues. TRPV1 agonists, including inflammatory lipid mediators, evoke membrane currents in isolated vascular myocytes to persistently constrict arteries, decrease coronary flow and increase blood pressure. Furthermore, these effects are retained after ablation of sensory nerves indicating an arteriole-mediated signaling mechanism. These data reveal a fundamental mechanism for transducing inflammatory stimuli into arterial constriction.

## Methods

### Animals

All experimental procedures involving mice and rats were approved by the Georgetown University and George Washington University Animal Care and Use Committee and the Ethics Committee on Animal Research of the University of Debrecen. Both Wistar and Sprague-rats (250-450 g) and C57Bl mice (25-30 g) were housed at 24-25°C and had *ad libitum* access to a standard laboratory chow and water.

### Mouse lines

The TRPV1-Cre transgenic mouse line (donated by Dr. Mark Hoon, NIH) was created using a BAC transgene containing the entire TRPV1 gene/promoter (50 kbp of upstream DNA) and IRES-Cre-recombinase.^21^ Importantly, Cre expression in this mouse faithfully corresponds with the expression of endogenous TRPV1. The TRPV1-Cre (hemizygous) mice were crossed with ai9 ROSA-stop-tdTomato mice (The Jackson Laboratory). The TRPV1^PLAP-nlacZ^ mice (Jackson Laboratory) were developed by Allan Bausbaum and colleagues (UCSF) to express human placental alkaline phosphatase (PLAP) and nuclear lacZ under the control of the TRPV1 promoter.^14^ The targeting construct contains an IRES-PLAP-IRES-nlacZ cassette immediately 3’ of the TRPV1 stop codon, which permits the transcription and translation of PLAP and nlacZ in cells expressing TRPV1 without disrupting the TRPV1 coding region. TRPV1-null mice were purchased from The Jackson Laboratory. TRPV1-Cre:ChR2/tdTomato mice were generated by crossing TRPV1-Cre mice with ChR2/tdTomato mice (The Jackson Laboratory).

### X-gal staining

TRPV1^PLAP-nlacZ^ mice were anesthetized with isoflurane and perfused through the heart using PBS (0.1 M, pH 7.3) followed by ice-cold 2-4% buffered paraformaldehyde. Whole skinned mice, brains, hearts, and trunk arteries were dissected and postfixed in 2-4% buffered paraformaldehyde on ice for 90 min, after which they were washed in PBS (containing 5 mM EGTA and 2 mM MgCl_2_) on ice and stained in X-gal solution (containing 1 mg/ml X-gal, 5 mM K_3_Fe(CN)_6_, 5 mM K_4_Fe(CN)_6_, 0.01% deoxycholate, and 2 mM MgCl_2_ in PBS) at 37°C overnight. nLacZ staining was imaged *in situ*, in heart sections (120-150 *µ*m thick) and in isolated arteries. To calculate the density of the signal, defined arteries and arterioles were isolated from TRPV1^PLAP-nlacZ^ and wild-type mice, stained and photographed in parallel. Densitometry was performed with ImageJ to yield density in arbitrary units (normalized to the wild-type signal). To map arterial/arteriole TRPV1 expression we compared the density of X-gal staining in main trunk arteries and tributaries, heart and brain. The distribution of intensities revealed 5 broad peaks (a baseline and four positive peaks, for example see **Fig. 4B**) that were color-coded from zero (dark blue) to a maximum (red).

**Figure 1.**
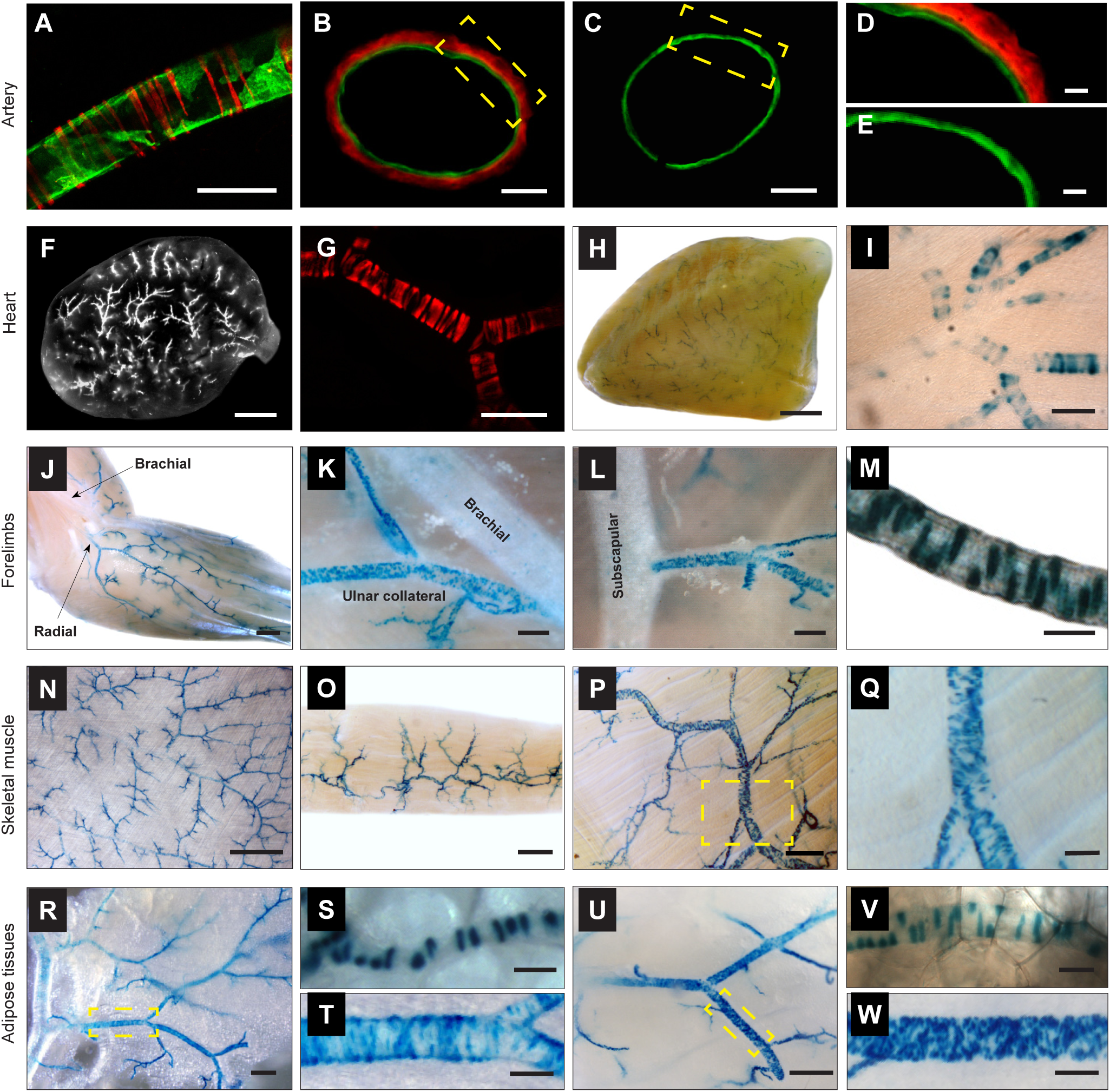
TRPV1 reporter mice reveal expression of TRPV1 in vascular smooth muscle of myocardium, skeletal muscle and fat. (**A**), TdTomato fluorescence in a muscle artery from a TRPV1-Cre:Tomato mouse. (**B, D**), Endothelium is stained with DioC18 (green). LacZ and CD31 immunostaining in artery cross-sections from a TRPV1^PLAP-nlacZ^ and (**C, E**), WT mouse. (**F, G**), Analysis of whole hearts and transverse heart sections from TRPV1-Cre:tdTomato or (**H, I**), TRPV1^PLAP-nlacZ^ mice reveals TRPV1 expression in small arterioles of the ventricular myocardium. (**J**-**M**), Nuclear LacZ staining in forelimb arteries, (**N**), latissimus dorsi, (**O**), gracilis and (**P, Q**), trapezius skeletal muscles, and arteries supplying (**R to T**), white and (**U to W**), brown adipose tissues. Insets (yellow boxes) in **P, R** and **U** are expanded in **Q, T** and **W** respectively. Scale bar: 1mm (F, H, J, N); 300 *µ*m (O, K, R, U); 100 *µ*m: (A, G, I, L, Q, T, W); 20 *µ*m (B, C, M, S, V); 10 *µ*m (D, E).

**Figure 2.**
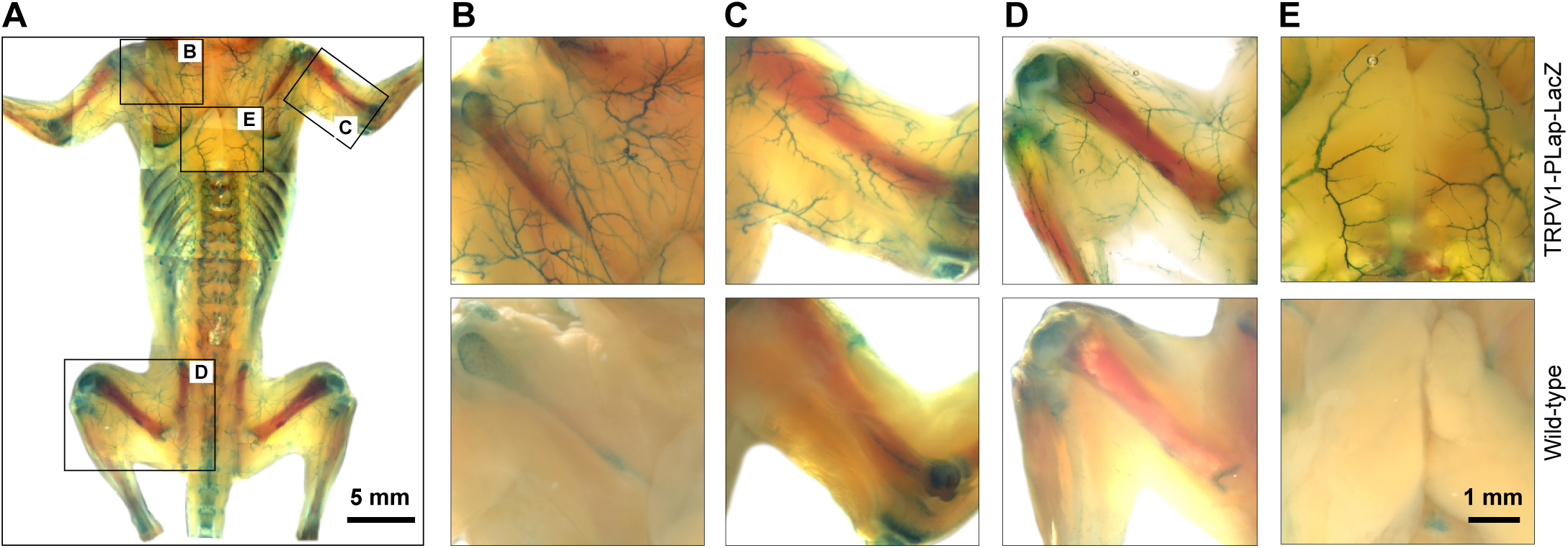
Arterial TRPV1 expression in a whole mouse preparation. (**A**-**D**), Whole-animal nLacZ staining in a 2 week old TRPV1^PLAP-nlacZ^ mouse (skin removed) versus a control (wild-type) mouse shows extensive arterial TRPV1 expression in skeletal muscles and (**E)**, interscapular brown adipose tissue. Note non-specific staining in bone tissues.

**Figure 3.**
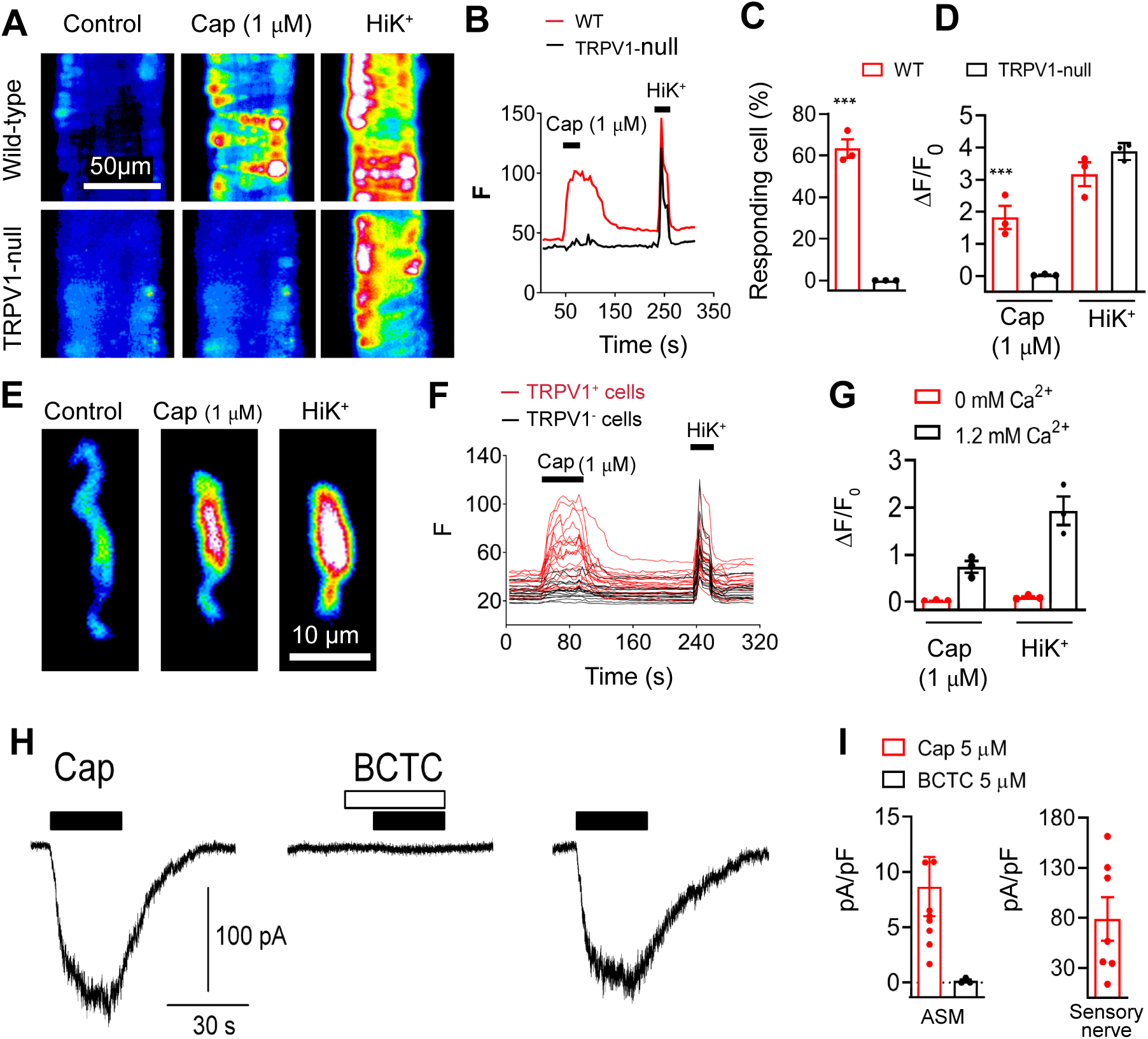
TRPV1 functionality in arterial smooth muscle cells. (**A-D)**, Capsaicin (1 μM) and KCl (50 mM) evoked Ca^2+^ signaling in isolated cerebellar arteries from wild type and TRPV1-null mice (n = 40 - 70 cells in the groups from 3 independent experiments, *\*\*\*P < 0.001*). (**E** and **F**), Capsaicin-evoked Ca^2+^ signaling in dissociated ASM cells from TRPV1-Cre:tdTomato mice is restricted to TRPV1^+^ cells (n = 20 - 25 cells per group). (**G**), Capsaicin and K^+^ evoked responses require extracellular Ca^2+^ (n = 20 – 30 cells per group). (**H**), Representative current traces in a voltage-clamped ASM cell (10 pF) in response to capsaicin (filled bars, 5 μM) with or without the TRPV1 antagonist BCTC (empty bars, 5 μM), and recovery after washout. (**I**), Mean current density in ASM cells for capsaicin (n = 9) and capsaicin + BCTC (n = 3), and in sensory nerves (n=7).

**Figure 4.**
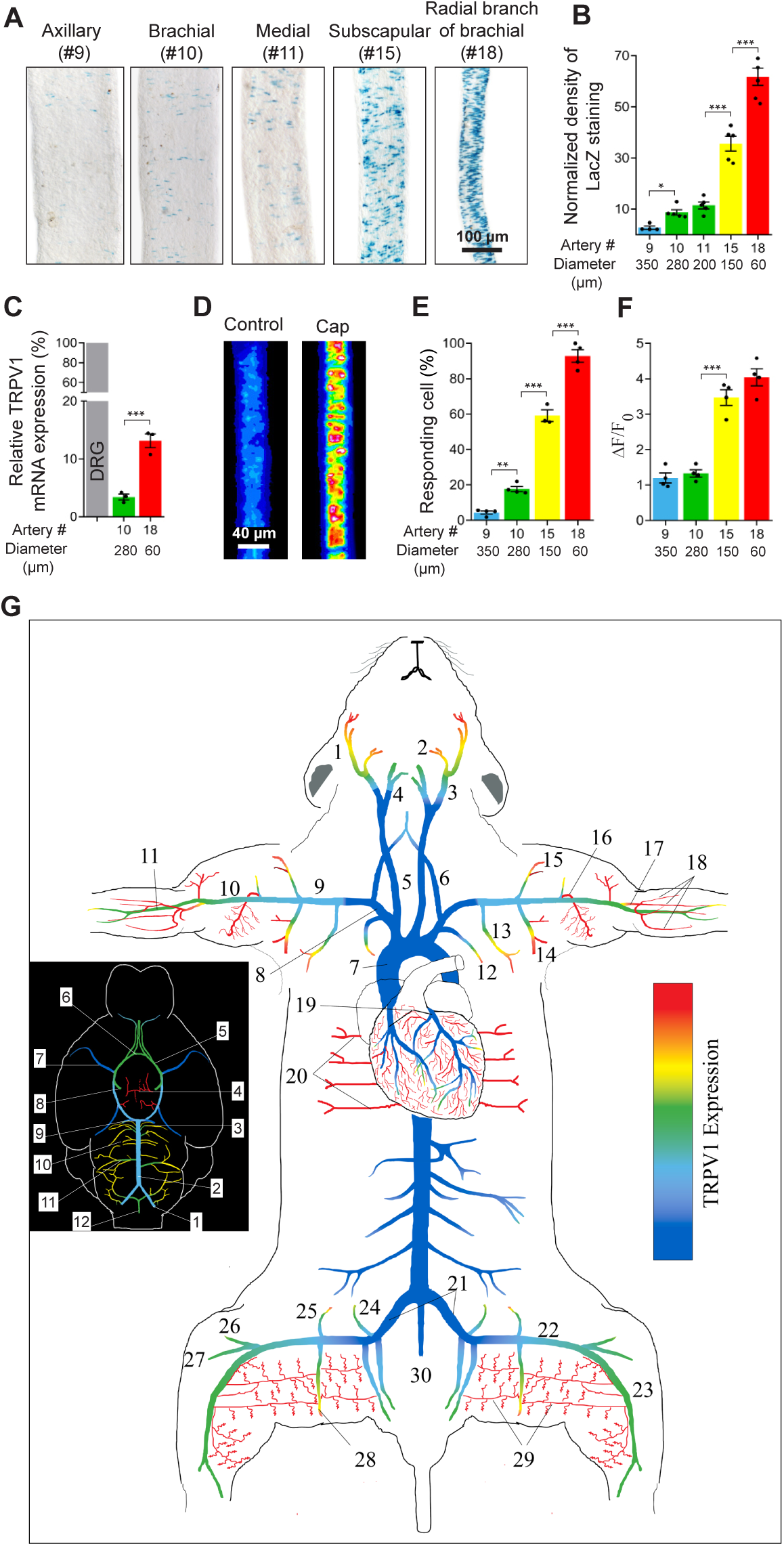
An arterial map of functional TRPV1 expression. (**A** and **B**), TRPV1 expression (nuclear LacZ staining) in forelimb arteries and muscle branches versus vessel diameter (n = 4 - 6 arteries from 5 mice per group, *\*P < 0.05, ***P < 0.001*). **(C)**, TRPV1 mRNA measured by qRT-PCR in small and large-diameter arteries relative to DRG (n = 3 - 4 mice), \*\*\**P* < 0.001. (**D to F**), Capsaicin (1 μM) evoked Ca^2+^ responses in forelimb arteries of different diameter (n ≥ 85 - 150 cells per group from 3 independent experiments, *\*\*P < 0.01, ***P < 0.001*). (**G**), Heat-map of TRPV1 expression in arteries based on TRPV1^PLAP-nlacZ^ mice (n ≥ 15 mice) and confirmed by functional imaging. Inset shows the density of TRPV1 expression in cerebral arteries. Arterial color-coding is similarly applied to ***B*** *to* ***F***. Artery nomenclature is included in table S1.

### TRPV1 mRNA analysis

RNA was extracted and purified using RNAqueous Micro Kit (Invitrogen). Quantitative PCR analysis was performed with *Taq*man Fast Advance Mastermix (Life Technologies), with TRPV1 gene probe labeled by FAM (Mm01246302_m1 from Life Technologies) and GAPDH control gene expression probe labeled by VIC (Mm99999915_g1). Thermal cycling of the PCR reaction was as follows: 50°C for 2 min, 95°C for 3 min, 50 cycles at 95°C for 15 s and 58°C for 1 min. Data were collected at the end of the 58°C anneal/extend step. Data were analyzed after the reaction using the Auto Ct function of the SDS 1.4 software (Life Technologies), and the reactions were considered to have passed quality control if the standard deviation of the Ct values were less than 0.5. The comparative Ct method was used to present gene expression of TRPV1 relative to GAPDH.

### Immunostaining and DiOC_18_ labeling

Mice were anesthetized with isoflurane and perfused through the heart using PBS (0.1 M, pH 7.3) followed by ice-cold 2-4% paraformaldehyde (in PBS). For endothelial labeling mice were perfused with the green fluorescent dye 3,3′-dioctadecyloxacarbocyanine perchlorate (DiOC18_3_; Sigma). DiOC18_3_ stock was prepared at 3 mg/ml in EtOH and diluted 50x in PBS. Arteries were isolated, fixed in 4% buffered paraformaldehyde for 1.5 h and stored in 30% sucrose overnight. Arteries were then embedded in low temperature agarose before sectioning (15 *µ*m). Sections were stained with primary antibodies, anti-LacZ (1:100, The Developmental Studies Hybridoma Bank, University of Iowa) and anti-CD31-FITC (1:50, Biolegend) followed by a goat anti-mouse IgG-AlexaFluor 555 secondary antibody (1:300, Invitrogen). Images were acquired by confocal microscopy.

### Sensory nerve culture

Dorsal root and nodose ganglia were obtained from adult mice (C57BL/6J) trimmed, digested with collagenase, and cultured in Neurobasal plus 2% B-27 medium (Invitrogen), 0.1% L-glutamine, and 1% penicillin/streptomycin on poly-D-lysine-coated glass coverslips at 37°C in 5% CO_2_. Neurons were used within 24 –36 h of culture.

### Arterial smooth muscle (ASM) cell isolation

Radial artery branch (artery #18, see **Fig. 4G**) and cerebellar branch (cerebral artery #3, see **Fig. 4G**) were washed in Mg^2+^-based physiological saline solution (Mg-PSS) containing 5 mM KCl, 140 mM NaCl, 2 mM MgCl_2_, 10 mM Hepes, and 10 mM glucose (pH 7.3). Arteries were initially digested in papain (0.6 mg/ml) (Worthington) and dithioerythritol (1 mg/ml) in Mg-PSS at 37°C for 15 min, followed by a 15-min incubation at 37°C in type II collagenase (1.0 mg/ml) (Worthington) in Mg-PSS. The digested arteries were washed three times in ice-cold Mg-PSS solution and incubated on ice for 30 min. After this incubation period, vessels were triturated to liberate smooth muscle cells and stored in ice-cold Mg-PSS before use. Smooth muscle cells adhered loosely to glass coverslips and were studied within 6 hours of isolation.

### Ca^2^ imaging

ASM cells and arteries were respectively loaded with 5 μM and 10 μM Fluo-4-AM (Invitrogen, Thermo Fisher Scientific) in a buffer solution containing 140 mM NaCl, 4 mM KCl, 1 mM MgCl_2_, 1.2 mM CaCl_2_, 10 mM HEPES, and 5 mM glucose (pH 7.3). Temperature was maintained at 32-35°C using a heated microscope stage (Tokai Hit). Bath temperature was verified by a thermistor probe (Warner instruments). ASM cells and arteries were imaged with 10X and 20X objectives using a Nikon TE2000 microscope with an excitation filter of 480 ± 15 nm and an emission filter of 535 ± 25 nm. The images were captured by a Retiga 3000 digital camera (QImaging) and analysis was performed offline using ImageJ.

### Electrophysiology

Whole-cell patch-clamp recordings were performed using an EPC8 amplifier (HEKA). Pipette resistances were in the range 3–4 MegaOhms and the current signal was low-pass filtered at 1–3 kHz and sampled at 4 kHz. The bath solution was the same as described for Ca^2+^ imaging (290 mosmol l^-1^). The pipette solution contained: 140 mM CsCl, 10 mM Hepes, 5 mM EGTA, and 1 mM MgCl_2_, pH 7.3.

### Ex vivo artery physiology

Skeletal muscle arteries (radial artery branch, artery #18, subscapular branch artery #14, gracilis artery #29, and cerebellar branch, cerebral artery #3, see **Fig. 4G**) were isolated and cannulated with glass micropipettes, and secured with monofilament threads. In some experiments arteries were denuded of endothelium by passing 1 ml of air followed by 1 ml of PSS through the lumen. Effective removal of the endothelium was confirmed by the absence of dilation of the arteries to ACh. The pipette and bathing PSS solution (containing 125 mM NaCl, 3 mM KCl, 26 mM NaHCO_3_, 1.25 mM NaH_2_PO_4_, 1 mM MgCl_2_, 4 mM D-glucose, and 2 mM CaCl_2,_) was aerated with a gas mixture consisting of 95% O_2_, 5% CO_2_ to maintain pH (pH 7.4). To maximally dilate arteries we perfused a PSS solution containing 0 CaCl_2,_ 0.4 mM EGTA and 100 *µ*M sodium nitroprusside (SNP). Arterioles were mounted in a single vessel chamber (Living Systems Instrumentation) and placed on a heated imaging stage (Tokai Hit), while intraluminal pressure was maintained by a Pressure Control Station (Stratagene) at 60 mmHg. Arteries were viewed with a 10X objective using a Nikon TE2000 microscope and recorded by a digital camera (Retiga 3000, QImaging). The arteriole diameter was measured at several locations along each arteriole using the NIH-ImageJ software’s edge-detection plug-in (Diameter) ^22^. The software automatically detects the distance between edges (by resampling a total of five times from adjacent pixels) yielding a continuous read-out ±SD of a vessel’s diameter.

Coronary arterioles were visualized in sagittal tissues slices (120-150 *µ*m) prepared from the hearts of TRPV1-Cre:tdTomato mice. Slices were perfused at room temperature with the same PSS solution as for pressurized arteries.

### Intravital imaging

Intravital imaging was performed in radial artery branches (about 60 *µ*m in diameter, artery #18 in **Fig 4G**). Animals were anesthetized with urethane (1.2 g/kg/IP). The forelimb was shaved and an incision was made. The skin and underlying muscle tissue were reflected to expose the brachial-radial artery junction. Both in WT and TRPV1-null mice, the arteries were visualized with a Zeiss stereomicroscope and illuminated with a low power blue light (using a standard GFP filter cube) exploiting the differential auto-fluorescence between tissue and blood. In TRPV1-Cre:ChR2/tdTomato mice, arteries were visualized with low power visible irradiation and stimulated with blue light. The exposed arteries were locally perfused (using a 250 *µ*m cannula connected to a valve-controlled gravity-fed perfusion system) with preheated buffer described for Ca^2+^ imaging. The surface tissue temperature (34-35°C) was measured via a thermistor (Warner Instruments) that was positioned next to the artery. Arteries were challenged with buffer without Ca^2+^ and with 1 mM EGTA to measure the passive diameter. The arteriole diameter was measured using ImageJ as described above for the *ex vivo* vessels.

### Coronary flow measurements

Sprague–Dawley rats (male, 300–350 g) were placed in a deep surgical plane of anesthesia by isoflurane inhalation, confirmed by lack of pedal reflex. The heart was then exposed via thoracotomy, quickly excised and rinsed in a bath of ice-cold perfusate. The aorta was rapidly cannulated then flushed with 500 units of heparin mixed with the perfusate that contained (in mM): 118 NaCl, 4.7 KCl, 1.25 CaCl_2_, 0.57 MgSO_4_, 1.17 KH_2_PO_4_, 25 NaHCO_3_, and 6 glucose. Hearts were then transferred to a retrograde perfusion system that delivered oxygenated (gassed with 95%O2-5%CO2) perfusate to the aorta at constant pressure (70mmHg) and 37±1°C. Coronary flow was measured using a tubing flowsensor (Transonic Systems) placed above the aortic cannula and was continuously acquired with the ECG using a PowerLab system (ADInstruments). Bolus injections of capsaicin were administered in-line above the aorta. BCTC was added to the perfusate reservoir. Data were analyzed off–line and the integral of coronary flow was calculated between the time of injection and the onset of the hyperemia response.

### Sensory nerve ablation

Neonate TRPV1^PLAP-nlacZ^ mice were treated with resiniferatoxin (50 ug/kg s.c.) at postnatal days 2 and 5. Sensory nerve ablation was confirmed at 8-12 weeks of age by LacZ staining of DRG ganglia. Newborn rats (at day 14 of life) were pretreated with Diaphyllin (Richter, Hungary), Bricanyl (Astra Zeneca, Hungary), and atropine (Egis, Hungary, 100 g/0,1 ml i.p.). Ten minutes later animals were injected with capsaicin (subcutaneously). The procedure was repeated for a total of five consecutive days. The total dose of capsaicin was 300 mg/kg, administered on a dose schedule of 10 mg/kg, 20 mg/kg; 50 mg/kg; 100 mg/kg; and 120 mg/kg on days 1 through 5, respectively. Rats were then kept in the animal facility for 10 weeks until experiments were performed.

### Measurement of capsaicin evoked sensory irritation

One drop (10 *µ*l) of capsaicin solution (50 μg/ml in physiological saline) was put into the right or left conjunctiva of the rat, in a random order. The number of eye wipes was counted during 60 s.

### Systemic blood pressure recording

The experiments were performed in anesthetized mice (urethane 1.2-1.5 g/kg/IP) and rats (thiopental 50 mg/kg/IP; supplemented by 5 mg/kg/IV if needed). After anesthesia, mice or rats underwent cannulation of the carotid artery and jugular vein as follows:

### Surgical preparation in the mouse

After the depth of anesthesia was confirmed by lack of pedal and corneal reflexes, mice were intubated via the trachea after tracheotomy to maintain an open airway and to institute artificial respiration when necessary. Next, the left carotid artery and the right jugular vein were cannulated with a Millar catheter (1F SPR-1000) and a polyethylene tubing (PE-10), respectively, for monitoring arterial blood pressure and for systemic (intravenous) infusion of drugs. To monitor heart rate, a three-point needle electrode-assembly representing Lead II of the electrocardiogram (ECG) was attached subcutaneously to the right and left forelimbs along with a reference electrode to the left hindlimb. Both the Millar catheter and the ECG assembly were coupled to a PowerLab data acquisition system (ADInstruments). Before vessel cannulation, the adjacent left cervical vagus was carefully isolated from the left carotid artery. Body temperature was monitored by a digital rectal thermometer and maintained at 37 ± 1°C with an infrared heat lamp.

### Conscious blood pressure recordings

RTX-treated mice (8-12 weeks) were surgically implanted with in-dwelling jugular catheters (Instech labs., USA) 48 h before BP recordings. BP was measured by tail-cuff plethysmography (Coda6, Kent Scientific, USA) performed before and immediately after infusion of drugs.

### Surgical preparation in the rat

Similar to the mouse, following intubation of the trachea, the left carotid artery, and jugular vein were cannulated with a polyethylene tubing (PE50) to monitor blood pressure and infuse drugs, respectively. Blood pressure (and ECG) was continuously recorded via a pressure transducer connected to the Haemosys hemodynamic system (Experimetria, Budapest, Hungary). The ECG was recorded from the extremities of the animal using hypodermic metal needles inserted subcutaneously in accordance with the Einthoven method (I, II, III leads). As in the mouse, heart rate was determined from lead II of the ECG recordings and body core temperature was maintained at 37±1 °C with a temperature controlled infrared heating lamp.

### Drug administration

Intravenous infusion of drugs was initiated only when a stable baseline of blood pressure and heart rate was present. This was also the case when drugs were re-administered. Final drug solutions contained: capsaicin (saline with 0.4% EtOH), LPA (saline with 0.6% EtOH).

### Chemicals

Capsaicin, resiniferatoxin and BCTC were purchased from Tocris Bioscience or Adooq Bioscience and stock solutions were prepared in EtOH at 1 M and 100 mM, respectively. LPA C18:1, were purchased from Cayman Chemical. Unless otherwise indicated, all other chemicals were obtained from Sigma–Aldrich.

### Statistical analysis

Data were analyzed using Prism (GraphPad Software, La Jolla, CA) and are expressed as means±SEM. Unless otherwise stated, statistical significance was evaluated using t-test and one-way ANOVA with treatment interactions assessed by Tukey’s *post hoc* multiple comparisons test.

## Results

### TRPV1 in arteries is restricted to vascular smooth muscle

Previous studies using antibodies have described TRPV1 expression in both vascular smooth muscle,^23, 24^ and endothelium.^17, 18^ However, the potential for non-specific labeling, well demonstrated for TRPV1 antibodies,^24, 25^ limits the interpretation of these data. To better define arterial expression of TRPV1, we exploited two validated mouse reporter lines. The first, TRPV1-Cre:tdTomato,^21^ generates a very sensitive fate map of TRPV1 expression. The second, TRPV1^PLAP-nlacZ, 14^ generates expression of human placental alkaline phosphatase (PLAP) and nuclear β-galactosidase (nLacZ) under the control of the endogenous TRPV1 promoter. Analysis of arterioles from TRPV1-Cre:tdTomato mice revealed robust tomato fluorescence in vascular smooth muscle that did not extend to the endothelium labeled with DioC18 (green, Fig. 1A). Similarly, co-labeling arteries from TRPV1^PLAP-nlacZ^ mice with antibodies to LacZ and the endothelial marker, CD31, revealed distinct, non-overlapping staining of smooth muscle and endothelium (Fig. 1, B to E). Furthermore, we did not detect any TRPV1 reporter staining in large arteries (Fig. S1 and S2). Thus, TRPV1 expression in murine arteries appears to be restricted to vascular smooth muscle.

### Arteriolar TRPV1 is enriched in skeletal muscle, heart and adipose tissues

Next, we mapped TRPV1 expression through the arterial network of the mouse. Remarkably, we found that TRPV1 was highly enriched in small (resistance) arterioles (<150 μm diameter) of the heart, skeletal muscle and adipose tissues. In the heart, large epicardial arteries were devoid of TRPV1 expression but strong tdTomato fluorescence (Fig. 1, F and G) and nLacZ staining (Fig. 1, H and I) emerged as these arteries penetrated and branched the myocardial wall. Similarly, TRPV1 expression, though mostly absent in aorta and large trunk arteries (Fig. S1), erupted as arteries branched to supply skeletal muscle (Fig. 1, J to M and Fig. S2). Indeed, analysis of skeletal muscle in whole animal (Fig. 2) or isolated tissue preparations (Fig. 1, N to Q and Fig. S1, S2) revealed an abundant network of TRPV1-expressing arteries. Note that LacZ is nuclear restricted and therefore staining excludes labeling of sensory nerve fibers. Additionally, LacZ uniformly stained vascular smooth muscle of arteries in both white and brown adipose tissue (Fig. 1, R to W). We also detected TRPV1 reporter expression in select parts of the cerebral circulation, including prominent labeling in the hypophyseal portal arteries and small-diameter branches of the basilar arteries (Fig. S3), and in very small mesenteric arteries (<50 μm, Fig. S1, Q to T). In addition, microvessels or “vasa vasorum” supplying the wall of large arteries highly expressed TRPV1 (Fig. S1, B). In contrast, we found very limited TRPV1 expression in arteries of other tissues examined, including skin, lung, kidney and liver (data not shown).

Next, to confirm that TRPV1 is functionally expressed, we performed Ca^2+^ imaging in isolated arterioles and arteriolar smooth muscle (ASM) cells. The TRPV1-specific agonist, capsaicin, increased Ca^2+^ signaling in a subset of arterioles isolated from wild-type mice, whereas we observed no responses in the same arterioles from TRPV1-null mice (Fig. 3, A to D). Furthermore, capsaicin sensitivity in ASM cells isolated from TRPV1-Cre:tdTomato mice was restricted to Tomato-positive cells (Fig. 3, E and F), and was abolished after removing external Ca^2+^ indicating an essential role for Ca^2+^ entry (Fig. 3G). Finally, capsaicin (5 μM) evoked inward currents in voltage-clamped isolated ASM cells (mean 8.7 ± 2.7 pA/pF, n = 9) that were fully prevented by the TRPV1 antagonist, BCTC (Fig. 3, H and I). The peak current density was ∼11% of that observed in cultured sensory neurons (Fig. 3, I).

To confirm the fidelity of the TRPV1 reporter, we compared both TRPV1 mRNA levels and the capsaicin sensitivity of arteries with differential reporter expression. TRPV1 LacZ reporter expression was inversely proportional to the arterial diameter (Fig. 4B), thus we isolated arteries at different positions along the axial-brachial trunk and branches (Fig. 4A). Quantitative PCR showed that TRPV1 mRNA levels were greater in smaller diameter arterioles peaking at ∼13% of DRG levels (Fig. 4C). Furthermore, results of Ca^2+^ imaging showed that the number of ASM cells responding to capsaicin (Fig. 4, D and E) and the magnitude of the Ca^2+^ signal (Fig. 4, D and F) increased in proportion to the TRPV1 reporter signal (nLacZ staining density). Based on TRPV1 reporter analysis, validated by functional imaging, we constructed a heat map of TRPV1 expression in the mouse arterial system (Fig. 4G). This map highlights the hierarchical distribution of TRPV1 becoming abundant in small-diameter resistance arterioles of skeletal muscle and heart (adipose is not represented here).

### TRPV1 constricts arterioles *ex vivo* and *in vivo*

To identify a physiological role for vascular TRPV1, we studied contractility in isolated, pressurized arteries. Capsaicin (1 μM) constricted (by ∼80-85%) arterioles isolated from wild-type (Fig. 5, A and B) and TRPV1-Cre:tdTomato mice (Fig. S4, A and B) without affecting vessels from TRPV1-null mice. Analysis of the response to varying capsaicin concentrations revealed a half-maximal effect at ∼150 nM that was unaffected by removing the vascular endothelium (Fig. 5C and Fig. S4, C to E), consistent with a selective action of capsaicin at arterial smooth muscle. To measure the *in vivo* functionality of arterial TRPV1, we performed intravital imaging of radial branch arteries. Local administration of capsaicin constricted these arteries (by ∼90%) without affecting nearby veins, or arteries in TRPV1-null mice (Fig. 5, D and E, Fig. S4 F and G, and Movies 1, 2). As an additional test of functional arterial TRPV1 expression, we generated mice expressing channelrhodopsin-2 (ChR2) under the control of TRPV1 gene regulatory elements (TRPV1-Cre:ChR2/tdTomato mice) to enable photo-control of cells expressing TRPV1. Control experiments revealed that expression of the TRPV1-Cre reporter and therefore ChR2, accurately reflect contemporaneous TRPV1 expression (Fig. S4A). Blue light rapidly and reversibly constricted arterioles in these animals, in both *ex vivo* and *in vivo* preparations, without affecting the caliber of the veins (Fig. 5E and Fig. S4, A and B) or arteries from TRPV1-Cre^-/-^ mice (data not shown). Thus, optogenetic sensitivity of arteries confirms TRPV1 expression in vascular smooth muscle, and can be exploited to control vessel diameter.

**Figure 5.**
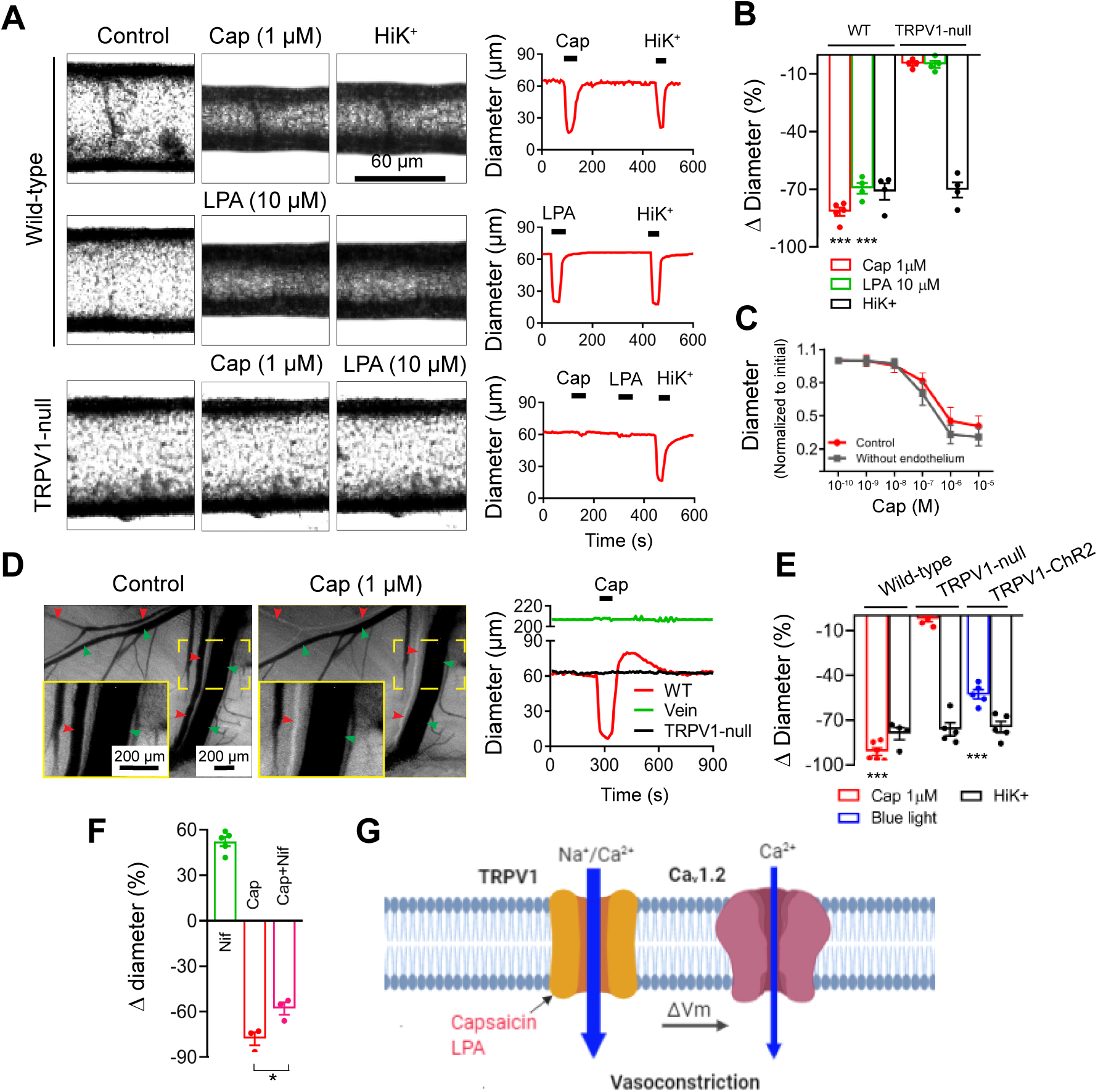
TRPV1 agonists constrict skeletal muscle arterioles. (**A, B** and **C**), Capsaicin (1 μM) and LPA (C18:1, 10 μM) constrict isolated, pressurized (60 mmHg) arteries from wild-type but not TRPV1-null mice. (**C**), Capsaicin constricts intact and endothelium denuded gracilis arteries with similar potency (n = 6). (**D** and **E**), Intravital imaging shows that capsaicin constricts radial branch arteries (red arrowheads) without affecting veins (green arrowheads). The inset (solid yellow) show expanded view of the designated area (broken yellow). Arteries from TRPV1-null mice are unresponsive to capsaicin and blue-light constricts arteries from TRPV1-Cre:ChR2 mice (n = 4 - 7 arteries from 3 - 5 mice per group, *\*\*\*P<0.001*). (**F**), *In vivo* arteriole diameter following treatment with nifidepine (3 μM), capsaicin, and nifedipine/capsaicin, (n = 3 – 5 arteries from 3 mice per group, *\*P < 0.05*). (**G**), Proposed model of Ca^2+^-entry pathways underlying capsaicin/LPA induced vasoconstriction.

Finally, we tested the relative contribution of Ca^2+^ entry via TRPV1 or voltage-gated channel channels. Indeed, the L-type voltage-gated Ca^2+^ channel (Cav1.2) is a major determinant of resting tone in arterioles. We found that nifedipine (3 μM) relaxed *in vivo* arterioles, but only partly (by ∼30%) inhibited the constriction evoked by a saturating concentration of capsaicin (Fig. 5F), indicating that Ca^2+^ influx through fully-activated TRPV1 channels is sufficient to constrict arteries, while L-type channels amplify the magnitude of the constriction. Figure 5G, summarizes the signaling pathways for TRPV1-mediated vasoconstriction; Ca^2+^ entry via TRPV1 and depolarization evoked Ca^2+^ entry via Cav1.2.

### TRPV1 constricts coronary arterioles and decreases coronary flow

The striking expression of TRPV1 in the coronary vasculature (Fig. 1) prompted us to explore functional effects of TRPV1 activation. Analysis of heart sections from TRPV1 reporter mice (Fig. 6, A and B) revealed that TRPV1 expression is restricted to small arterioles that branch from the large coronary arteries (yellow arrowheads). Application of capsaicin 1 μM to sagittal slice preparations (∼150 μm) of living heart tissue constricted these small arterioles and the magnitude of the response was inversely proportional to artery diameter (Fig. 6, C to E). Next, we examined a role for TRPV1 in the regulation of coronary flow in isolated rat hearts. Administration of capsaicin (by 10 s in-line infusion) decreased flow in a dose-dependent manner, followed by a rebound hyperemia (Fig. 6, F to H). The decrease in flow occurred without a change in heart rate and was fully prevented by pretreatment with BCTC.

**Figure 6.**
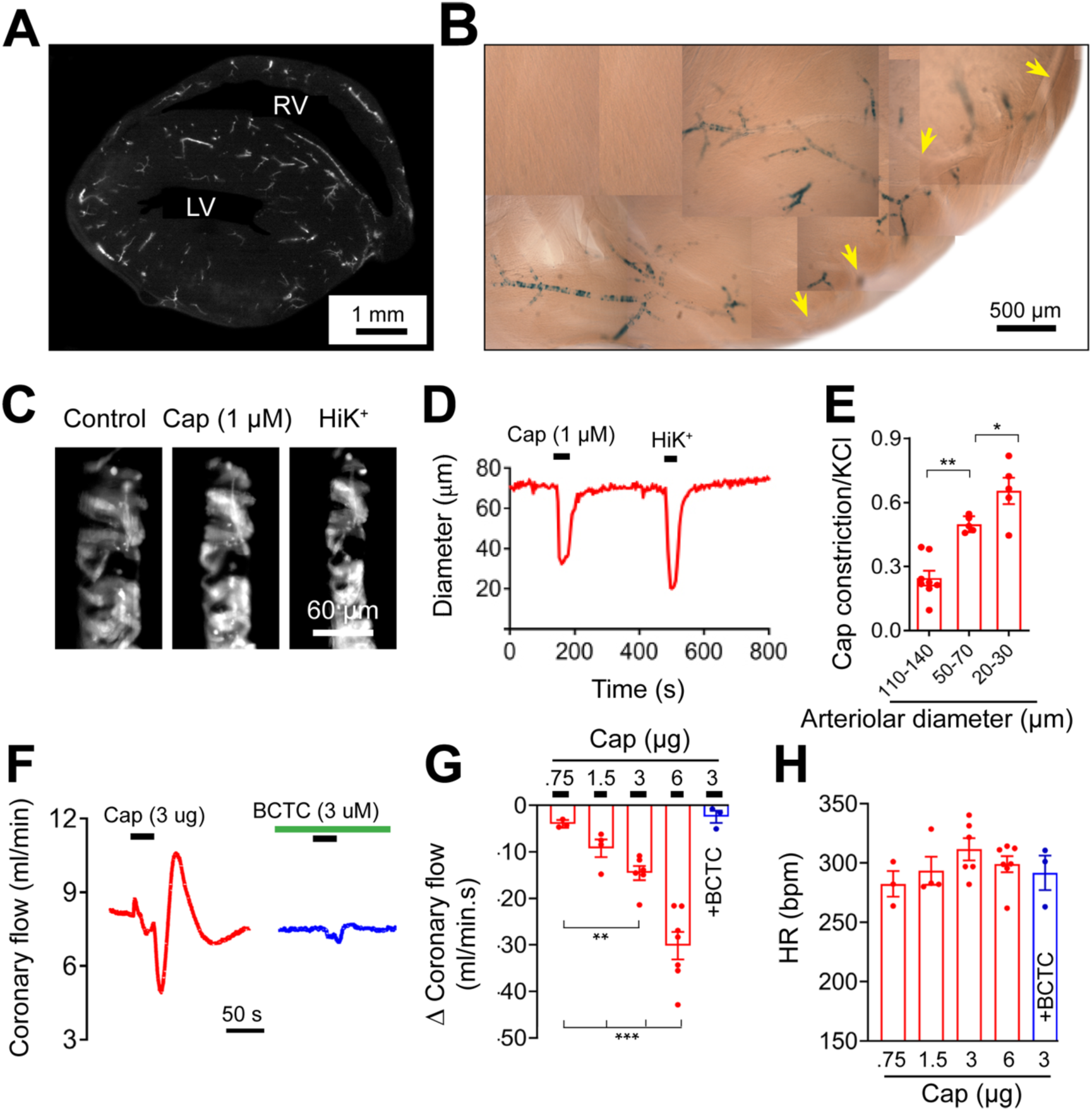
Capsaicin constricts coronary arterioles and reduces coronary perfusion. (**A** and **B)**, Sagittal sections (150 μm) of hearts from TRPV1-Cre:tdTomato and TRPV1-nLacZ mice reveal TRPV1 expression in small arteriole branches (yellow arrowheads denote the left coronary artery). (**C-E)** Capsaicin preferentially constricts small arterioles *in situ* in slice preparations from hearts of TRPV1-Cre:tdTomato mice. Data are normalized to KCl (40 mM) (n =5). **(F-H)**, Capsaicin dose-dependently decreases coronary perfusion in isolated rat hearts without affecting the heart rate. BCTC abolishes the effect of capsaicin (n = 3-7, \**P<0.05, **P<0.01, ***P<0.001*).

### Arterial TRPV1 regulates systemic blood pressure

The extensive expression in arteriolar smooth muscle makes TRPV1 well situated to influence systemic blood pressure (BP). We therefore tested whether TRPV1 agonists would alter BP as predicted by their profound vasoconstrictive effects detailed above. Indeed, intravenous (IV) administration of capsaicin to anesthetized mice markedly increased BP, while TRPV1-null mice exhibited no responses to capsaicin demonstrating a selective action at TRPV1 (Fig. 7, A and B and Fig. S5, A to C). Further, we observed equivalent BP responses to capsaicin in conscious mice (Fig. 5I and Fig. S5H) ruling out side effects of anesthesia. Similarly, in rats, capsaicin produced a dose-dependent increase in BP with an approximate 60 mmHg rise in systolic and diastolic blood pressure observed at the highest dose tested, (Fig. 7C, Fig. S5, K to M). The peak responses to capsaicin occurred without any significant changes in heart rate (Fig. S5, D, G, I, N and P), demonstrating a predominant effect on peripheral vascular resistance.

**Figure 7.**
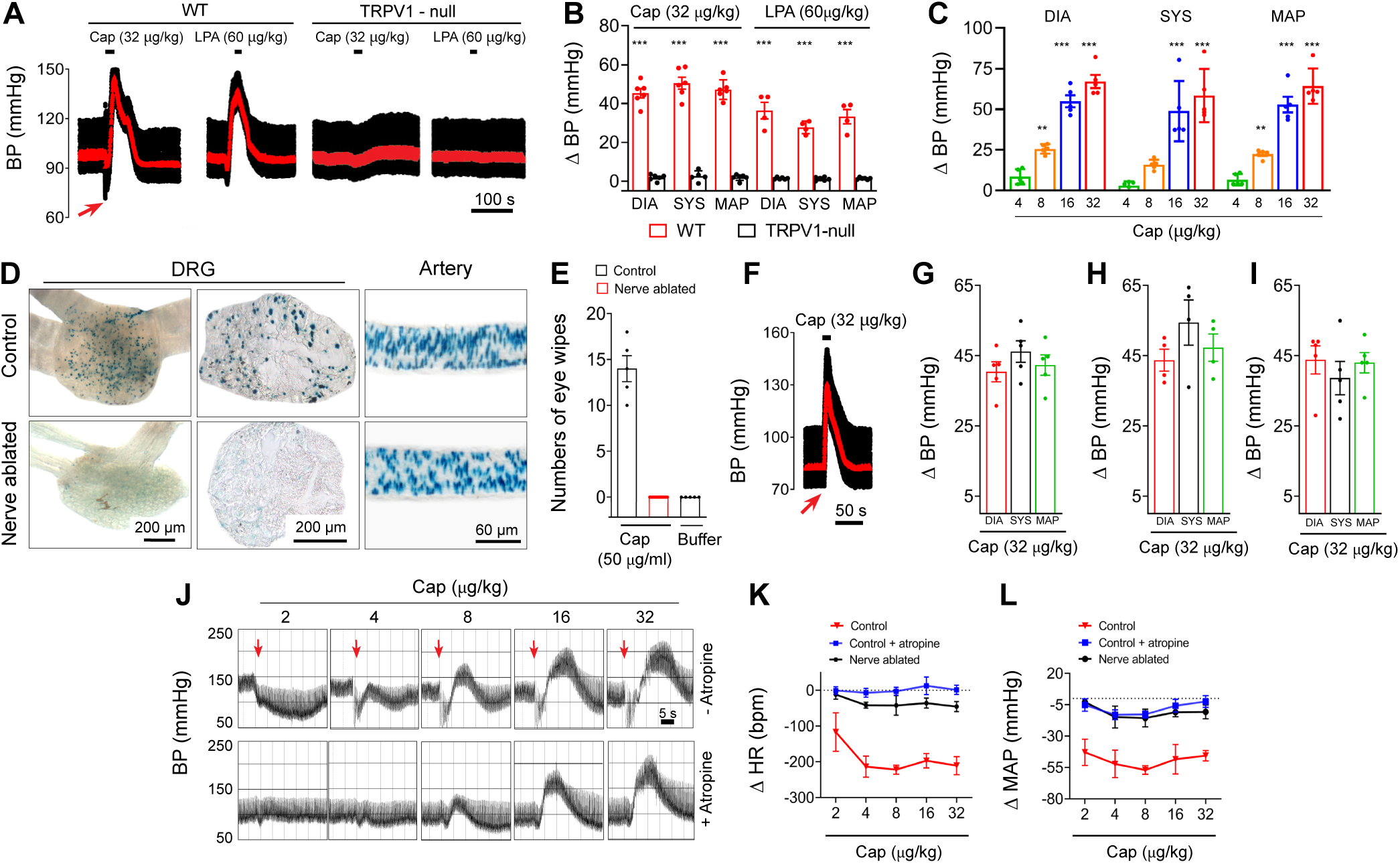
TRPV1 regulates systemic blood pressure. (**A** and **B**), Blood pressure (BP) changes in wild-type and TRPV1-null mice in response to intravenous (IV) infusion (20 s) of capsaicin or LPA (C18:1)(n = 4 - 7, *\*\*\*P < 0.001*). Mean arterial pressure is shown in red. (**C**), Mean changes in systolic BP, diastolic BP and MAP in rats during bolus IV capsaicin (n = 6, *\*\* P < 0.01, ***P < 0.001*). (**D**), nuclear LacZ staining in L5 dorsal root ganglion (section and whole ganglion) and arteries from mice with or without neonatal RTX treatment. (**E**), Mean eye-wipes in response to capsaicin in nerve-ablated rats (n = 5). (**F** and **G**), BP responses to IV capsaicin in sensory nerve ablated mice (n = 4) and (**H**), rats (n = 5). (**I**), Change in BP in conscious mice in response to IV administration (20 s) of capsaicin (n = 4). (**J**), BP recordings in a rat in response to escalating bolus IV doses of capsaicin with or without atropine pretreatment [Note: atropine abolishes the rapid decrease in BP reflecting a Bezold-Jarisch reflex]. (**K** and **L**), Mean changes in heart rate and MAP measured immediately after bolus IV capsaicin in rats (n = 6).

Although the pressor response to capsaicin is consistent with actions at arterial TRPV1, these data do not exclude a contribution of TRPV1 in perivascular sensory nerves. TRPV1-expressing nerves may affect blood flow by releasing vasoactive peptides such as CGRP,^26^ or neurokinins. ^19, 20^ Therefore, to discriminate between an arterial and neurogenic locus of TRPV1 signaling, we performed selective ablation of TRPV1-expressing sensory nerves. Resiniferatoxin or capsaicin administered systemically to neonates causes permanent deletion of most TRPV1-positive sensory nerves.^27, 28^ In contrast, TRPV1-expressing arterial smooth muscle exhibits full functional recovery from this treatment,^29^ reflecting differences between neurons and vascular smooth muscle cells. Indeed, 8 weeks following RTX administration to TRPV1-nLacZ mice, we observed an almost complete loss of nLacZ staining in DRG neurons, whereas arterial staining was unaffected (Fig. 7D and Fig. S5E). Similarly, treating neonatal rats with capsaicin abolished subsequent nocifensive responses (eye-wipes) to capsaicin (Fig. 7E) consistent with ablation of TRPV1 expressing sensory neurons. Notably, administration of capsaicin to these sensory-nerve ablated mice (Fig. 7G and Fig. S5, F and H) and rats (Fig. 7H and Fig. S5O) evoked pressor responses similar to untreated controls. Thus, we conclude that sensory nerves play little to no role in the capsaicin-evoked BP rise.

In previous studies, capsaicin was shown to elicit a cardiopulmonary Bezold-Jarisch reflex consisting of a transient drop in BP, bradycardia and apnea.^30^ Indeed, we observed that capsaicin evoked a fast, transient depressor response that preceded the rise in BP (see arrowhead in Fig. 7A). At doses <4 μg/kg, bolus capsaicin only produced this transient depressor response accompanied by a decrease in heart rate (Fig. 7, J to L). At higher doses (from 8 μg/kg capsaicin), an additional pressor response emerged (Fig. 7J). Notably, pretreatment with atropine or sensory nerve ablation eliminated this transient depressor response to capsaicin, consistent with it being mediated by the vagus nerve, and unmasked the arteriolar TRPV1 mediated increase in BP.

### Lysophosphatidic acid constricts arterioles and increases blood pressure via TRPV1

Next, we tested the potential physiological function of vascular TRPV1. We hypothesized that the vasoconstrictive effect of some endogenous bioactive lipids is mediated by TRPV1 activation. We examined lysophosphatidic acids (LPA), a vasoconstrictor lipid species generated by platelets and atherogenic plaques,^31, 32^ potentially activating TRPV1.^33^ Notably, LPA species containing an unsaturated acyl chain are potent vasoconstrictors but whether cognate GPCRs for LPA (LPA_1-6_) mediate these effects is uncertain.^31, 34^ We found that LPA (C18:1) constricted skeletal muscle arterioles isolated from WT but not from TRPV1-null mice confirming a selective action at TRPV1 (Fig. 5, A and B). Furthermore, systemic administration of LPA (60 μg/kg) triggered a Bezold-Jarisch reflex and an increase in BP in a TRPV1-dependent manner (Fig. 7, A and B and Fig. S5, A to C). These data agree with previous observations that LPA exerts a pressor effect in several mammals (cats, rats, and guinea pigs) but notably not in rabbits, ^35, 36^ that possess a hypofunctional TRPV1 channel.^37^ In summary, our data show that TRPV1 contributes to the regulation of arteriolar tone and systemic BP by mediating vasoconstrictive responses to the inflammatory lipid LPA.

### Vascular TRPV1 is resistant to desensitization and mediates a persistent increase in blood pressure

In sensory nerves, TRPV1 ordinarily exhibits pronounced desensitization to agonists, reflected by a diminishing current response to repeated (tachyphylaxis) or sustained agonist application. ^38, 39^ Indeed, in voltage-clamped sensory neurons we found that the response to these forms of capsaicin application declined rapidly to less than 20% of the initial current (Fig. 8, A and C). In contrast, in isolated arterial smooth muscle cells, capsaicin evoked non-declining currents; the current response to the fourth application was 110% of the first application and there was no evident current decay during sustained treatment (Fig. 8, B and C). Similarly, repetitive or prolonged (5 - 20 minutes) systemic administration of capsaicin to mice evoked reproducible and sustained increases in BP (Fig. 8, D to G and Fig. S6). These pressor effects were unaffected by ablation of sensory nerves (Fig. 8G) and occurred without changes in heart rate upon the elevated BP (Fig. 8 E, F and I), reflecting a primary action of TRPV1 located in vascular myocytes. Finally, we tested whether LPA could generate persistent BP responses. Similar to capsaicin, slow infusion of LPA (C18:1, 75 μg/kg/min) produced sustained increases in BP without affecting heart rate (Fig. 8 F, H, and I).

**Figure 8.**
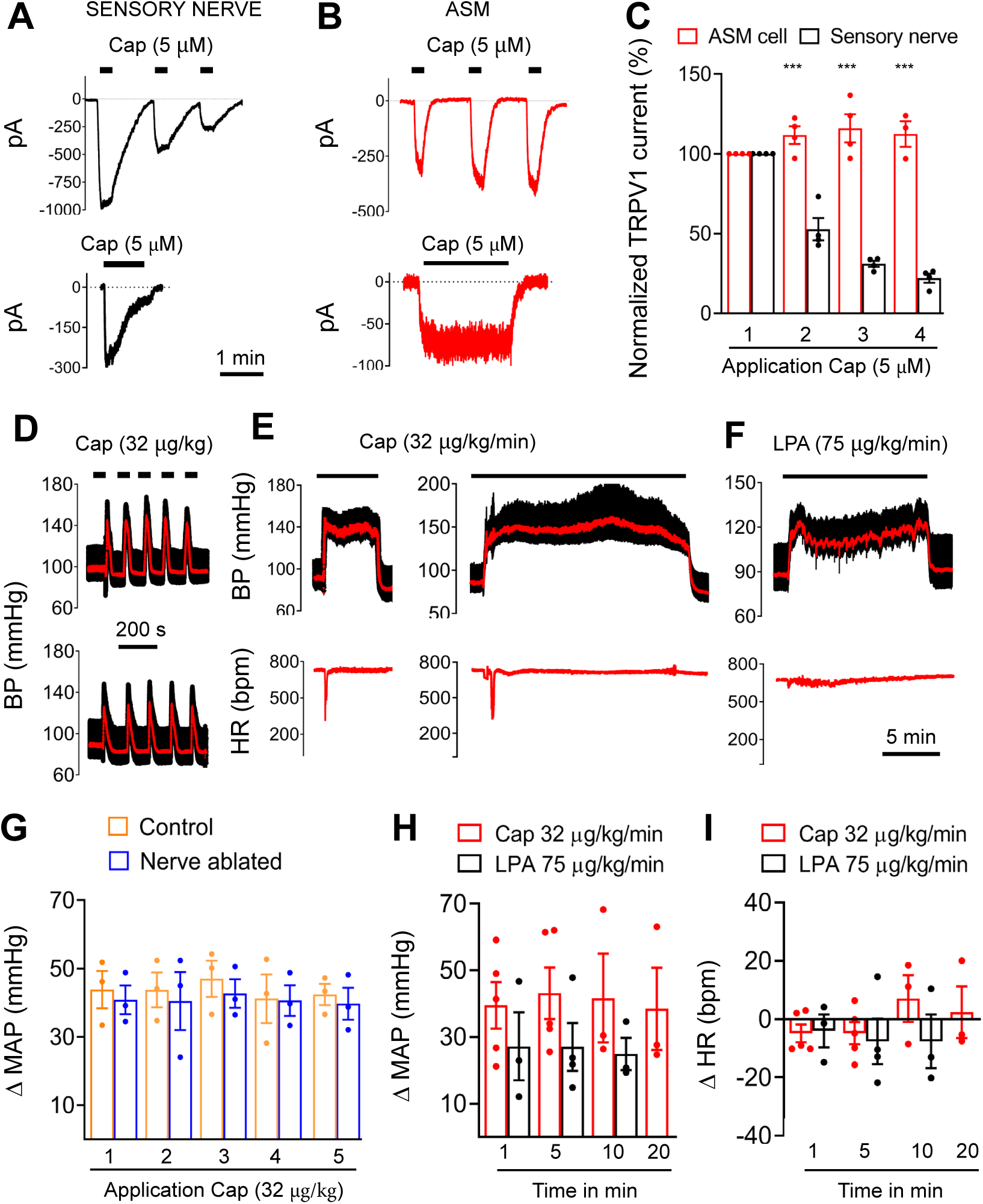
TRPV1 mediates persistent vasoconstriction. (**A** and **B**), Representative current traces in sensory neurons and ASM cells in response to repetitive or sustained application of capsaicin. (**C**), Mean normalized peak current evoked by repeated capsaicin treatment (n = 4), *\*\*\*P<0.01* (neurons versus ASM cell). (**D-F**), Representative BP traces in response to bolus or continuous infusion of capsaicin or LPA. (**G-I**), Mean changes in BP and heart rate (HR) in response to repeated or continuous capsaicin/LPA administration, n = 3 - 5.

## Discussion

TRPV1 is an ion channel with important roles in somatosensory transduction.^2, 12^ Our data reveal extensive expression of TRPV1 in the arterial circulation. Using combined molecular and functional analyses we found that TRPV1 localizes to the smooth muscle of arterioles supplying skeletal muscle, heart and adipose tissues. Notably, we did not detect TRPV1 in vascular endothelium and removal of endothelium had no effect on the efficacy/potency for capsaicin to constrict arteries. Furthermore, our analysis shows that TRPV1 predominates in small-diameter (<150 μm) arteries and is practically absent in large vessels. Quantitative mRNA measurements showed that TRPV1 expression in small arterioles is ∼13% of DRG levels. Similarly, the peak capsaicin-evoked current density in arterial smooth muscle cells was ∼11% of that recorded in sensory neurons. Several pertinent observations can be made about TRPV1 expression in vascular smooth muscle. First, given the extensive network of arteries in muscle and adipose tissues then the overall level of TRPV1 protein in the vasculature likely exceeds that in nerves. Second, activation of TRPV1 even at low levels, was nonetheless capable of markedly constricting skeletal muscle arteries, both *ex vivo* and *in vivo*, and coronary arteries leading to reduced coronary perfusion. The L-type Ca^2+^ channel (CaV1.2) blocker, nifedipine, only partly inhibited the response to a saturating concentration of capsaicin, indicating that Ca^2+^ entry via maximally activated TRPV1 *per se* can support constriction while depolarization induced activation of CaV1.2 amplifies the response. The amplification may become prominent during submaximal activation of TRPV1.

We found that systemic administration of TRPV1 agonists elicited large increases in BP. This pressor effect occurred without significant changes in heart rate and is consistent with a vasoconstrictor action of TRPV1. Although TRPV1-expressing sensory nerves, through the release of vasoactive peptides, could contribute to changes in BP, we found the pressor response was independent of neurogenic regulation. First, we studied mice and rats treated after birth with RTX or capsaicin to ablate the TRPV1–positive sensory nerve population. TRPV1 in the vasculature of these animals exemplified complete recovery when measured at 8-10 weeks, reflecting lower toxicity and /or replacement of nascent arterial smooth muscle cells. Notably, the BP response to capsaicin in these nerve-ablated animals was unchanged compared with control animals indicating that sensory nerves do not significantly contribute to TRPV1-mediated BP regulation. One exception to this result was the presence of a fast depressor response to capsaicin, most evident upon bolus administration. This BP decrease, accompanied by decreased heart rate and apnea, likely reflects the Bezold-Jarisch reflex,^30^ and was abolished after sensory nerve ablation or after atropine treatment to inhibit vagal cholinergic responses. Second, we found that TRPV1 channels in ASM cells resist desensitization in response to capsaicin stimulation. Significantly, this effect was recapitulated in systemic BP recordings where infusions of TRPV1 agonists produced sustained increases in BP without desensitization. Furthermore, during repetitive capsaicin treatment we observed that the Bezold-Jarisch reflex was evident only upon the first administration consistent with a pronounced desensitization of sensory nerves. Thus taken together, the sustained increases in BP correlate with the persistent opening of TRPV1 channels in ASM cells and not with the transient TRPV1 response in sensory nerves. Our finding that sensory nerves play little role in the persistent capsaicin regulation of BP may seem surprising, but is consistent with the plethora of earlier studies showing that sensory nerves primarily trigger dilation rather than constriction of arterioles, and that this effect is especially evident in the skin and other epithelial tissues.^19, 20^ Similarly, capsaicin triggers marked plasma extravasation (measured by Evans Blue) that is restricted to the skin, airways, and urogenital organs, that have prominent perivascular sensory nerves, and neurogenic inflammation is absent throughout the remainder of the circulation including brain, skeletal muscle and the heart.^19, 40^ Thus, any neurogenic vasodilatory action in response to systemic capsaicin would be swamped by the direct vasoconstrictor effects in arterioles.

What processes allow vascular smooth muscle TRPV1 to resist desensitization? Desensitization of TRPV1 is both Ca^2+^ and state dependent. In sensory nerves the removal of external Ca^2+^ ions abolishes capsaicin-induced desensitization consistent with a key role for Ca^2+^ signaling.^38, 39^ Direct binding of calmodulin,^41^ Ca^2+^-induced dephosphorylation,^42, 43^ or Ca^2+^-dependent hydrolysis of phosphatidylinositol 4,5-bisphosphate (PIP_2_),^44, 45^ have all been proposed as potential mechanisms. Structural changes in the TRPV1 channel may underlie desensitization. Evidence in support of this is provided by studies of the ultrapotent agonist, RTX, and double-knot spider toxin (DkTx) that do not induce desensitization even in the presence of external Ca^2+^. The results of cryo-EM analysis, ^4^ suggests that binding of RTX/DkTx stabilizes displacement of the pore helix which is a mobile element in gating, to produce sustained channel opening. In contrast, capsaicin does not stabilize movement of the pore helix, but instead engages the lower channel gate,^4^ leading to the appearance of flickery channel openings,^46^ that may facilitate the transition to desensitized states. Thus, in arteries resistance to capsaicin-induced desensitization may arise from modifications to the TRPV1 channel protein, or alternatively, the existence of different Ca^2+^ signaling pathways, but further studies are needed to understand the precise underlying mechanisms.

TRPV1 plays a critical role in pain signaling; diverse inflammatory mediators activate or sensitize TRPV1 located in sensory nerves to enhance nociception.^1, 2, 47^ Our data reveal that TRPV1 agonists, including capsaicin and LPA, act on arterial TRPV1 to mediate vasoconstriction and a sustained increase in systemic BP. Notably, LPA is produced by platelets and artherogenic plaques,^31, 32^ and is elevated in acute coronary syndrome,^48, 49^ associated with vasospasm of small coronary arteries. TRPV1 in the coronary microcirculation may therefore represent a prime target for mediating this vasoconstriction. Further, autotaxin, the rate-limiting enzyme for LPA production, is secreted abundantly by adipocytes,^50^ bringing the synthesis of LPA in close proximity to adipose arteries that highly express TRPV1. Notably, several lipoxygenase-dependent metabolites of arachidonic acid are potent TRPV1 agonists, including 12- and 15-(*S*)-HPETE, 5-(*S*)- and 5-(*R*)-hydroxyeicosatetraenoic acid (5-(*S*)-HETE and 5-(*R*)-HETE), 12-(*S*)-HETE, and leukotriene B_4_, ^51^ and may therefore be capable of constricting TRPV1-expressing arteries. Genetic studies also support a fundamental role for TRPV1 in the regulation of blood flow. Indeed, disrupting TRPV1 gene expression exacerbates ischemic reperfusion injury in the heart.^52^ Furthermore, in experimentally induced sepsis, TRPV1-null mice exhibit both a greater fall in BP and higher mortality than their wild-type counterparts,^53-56^ suggesting that TRPV1-mediated vasoconstriction during inflammation contributes to the homeostatic regulation of BP. Collectively, these observations indicate potentially important roles for TRPV1 in regulating vasoconstriction and BP in disease and injury states.

## Acknowledgments

We thank R. Gillis and R. Miyares for comments on the manuscript.

## Author contributions

T.P designed and performed most of the experiments, analyzed the data and helped write the manuscript; G.A. conceived and designed experiments, performed electrophysiology and wrote the manuscript; H.T. performed experiments; N.S. assisted with mouse BP measurements and helped write the manuscript; H.G and R.B. performed rat physiology and helped write the manuscript. A.T. helped design experiments and helped write the manuscript. R.R. and M.K. performed coronary flow experiments.

