## Supplemental Figures for "TRPV1 expressed throughout the arterial circulation enables inflammatory vasoconstriction"

| | | |
|---|---|---|
| (1) Superficial temporal | (16) Profunda brachii | ***Arteries in Inset*** |
| (2) Facial | (17) Ulnar collateral | (1) Vertebral |
| (3) External carotid | (18) Radial branches | (2) Basilar |
| (4) Internal carotid | (19) Coronary | (3) Superior cerebellar |
| (5) Common carotid | (20) Intercostal | (4) Hypophyseal portal |
| (6) Vertebral | (21) Common iliac | (5) Anterior cerebral |
| (7) Aortic arch | (22) Femoral | (6) Anterior communicating |
| (8) Subclavian | (23) Saphenous | (7) Middle cerebral |
| (9) Axillary | (24) Iliaco-femoral | (8) Internal carotid |
| (10) Brachial | (25) Superficial caudal epigastric | (9) Posterior cerebral |
| (11) Medial | (26) Medial proximal genicular | (10) Pontine |
| (12) Radial | (27) Popliteal | (11) Anterior inferior cerebellar |
| (13) Internal mammary | (28) Proximal caudal femoral | (12) Anterior spinal |
| (14) Lateral thoracic | (29) Gracilis | |
| (15) Subscapular | (30) Median coccygeal | |

***Supplementary Table 1.*** **Artery nomenclature for Figure 4.**

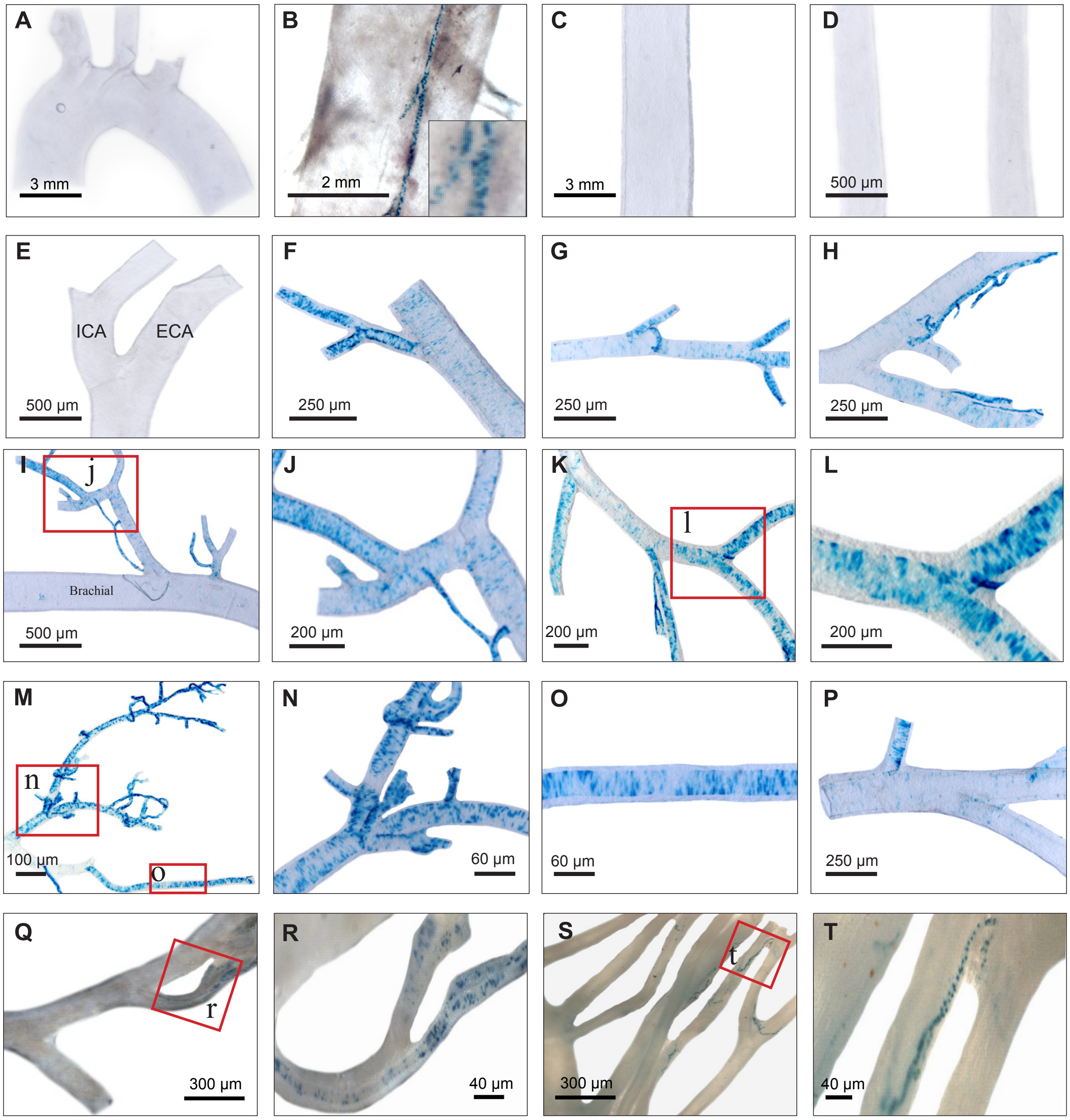


***Figure S1.*** **Restricted TRPV1 expression in isolated aorta, skeletal muscle and mesenteric arteries**. (**A**), nLacZ staining in the aortic arch and (**B**), descending aorta (showing TRPV1 expression is restricted to small feeding arteries "vasa vasorum", see inset) and (**C**), abdominal aorta and (**D**), common carotid and (**E**), external (ECA) and internal carotid (ICA) arteries and (**F**)**,** facial artery and (**G**), maxillary artery, and (**H**), superficial temporal artery and (**I, J**), axillary artery and ulnar collateral artery and (**K, L**),subscapular artery and branches and (**M-O**), muscle branches of brachial artery and (**P**), medial artery and branches and (**Q-T**), mesenteric arteries.

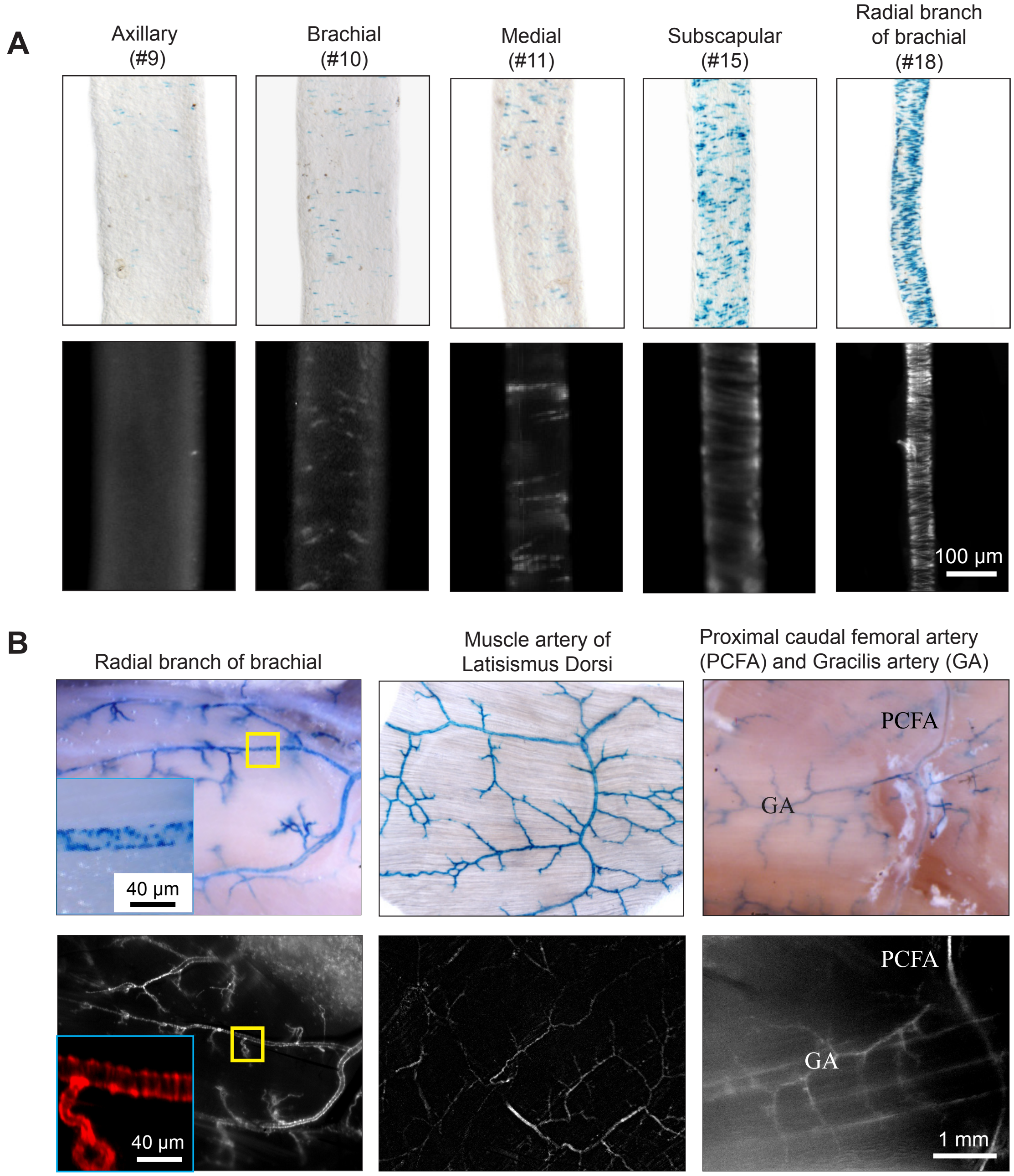


***Figure S2.*** **TRPV1 expression in arteries of forelimb and skeletal muscle.** (**A**) nLacZ staining (TRPV1$^{PLAP\text{-}nlacZ}$ mouse) and tdTomato fluorescence (TRPV1-Cre:tdTomato mouse) in forelimb trunk arteries, and (**B)** skeletal muscle arteries**.** Note that TRPV1 expression increases inversely proportional to arterial diameter (*A*) and with branching into tissue (*B*) and the correspondence of the nLacz staining and tdTomato signals.

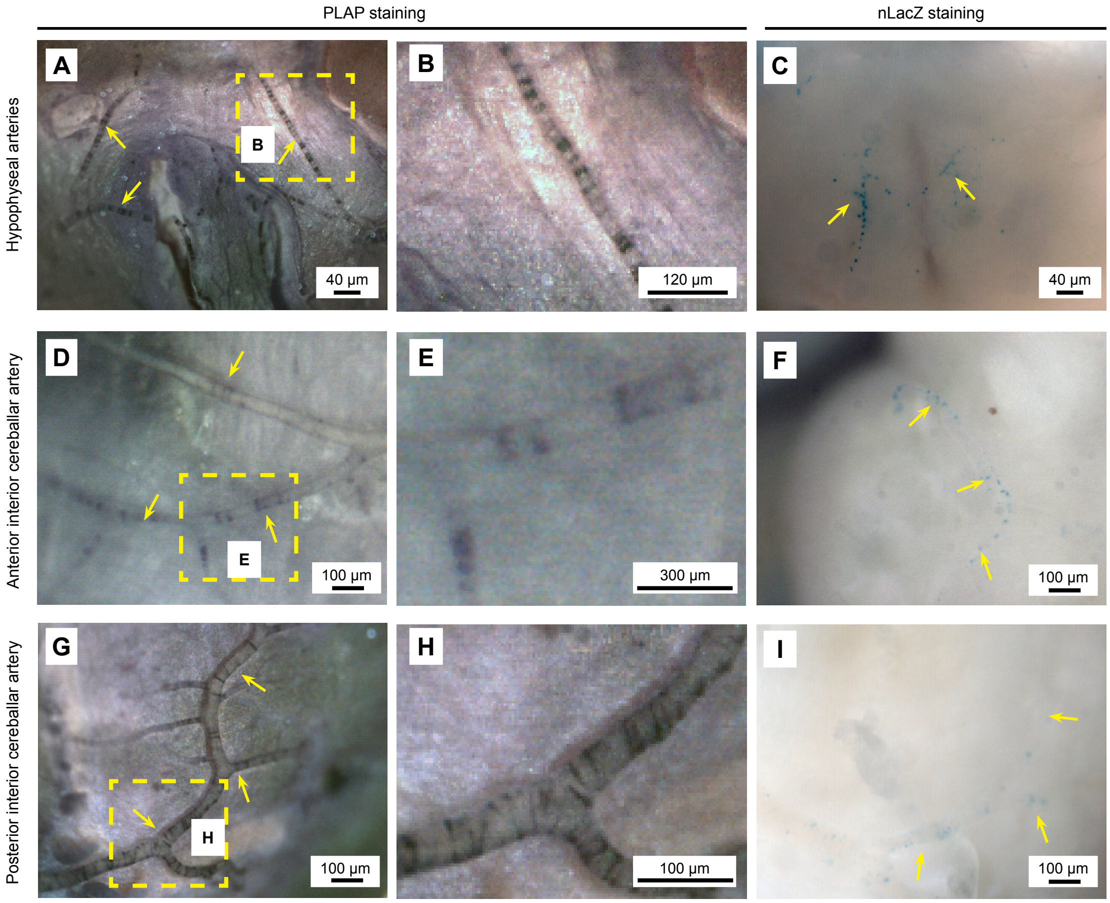


***Figure. S3.*** **TRPV1 expression in cerebral arteries.** PLAP and nLacZ staining in (**A** to **C**), hypophyseal portal arteries, (**D** to **F)**, anterior inferior cerebellar artery, and (**G** to **I )**, posterior inferior cerebellar artery.

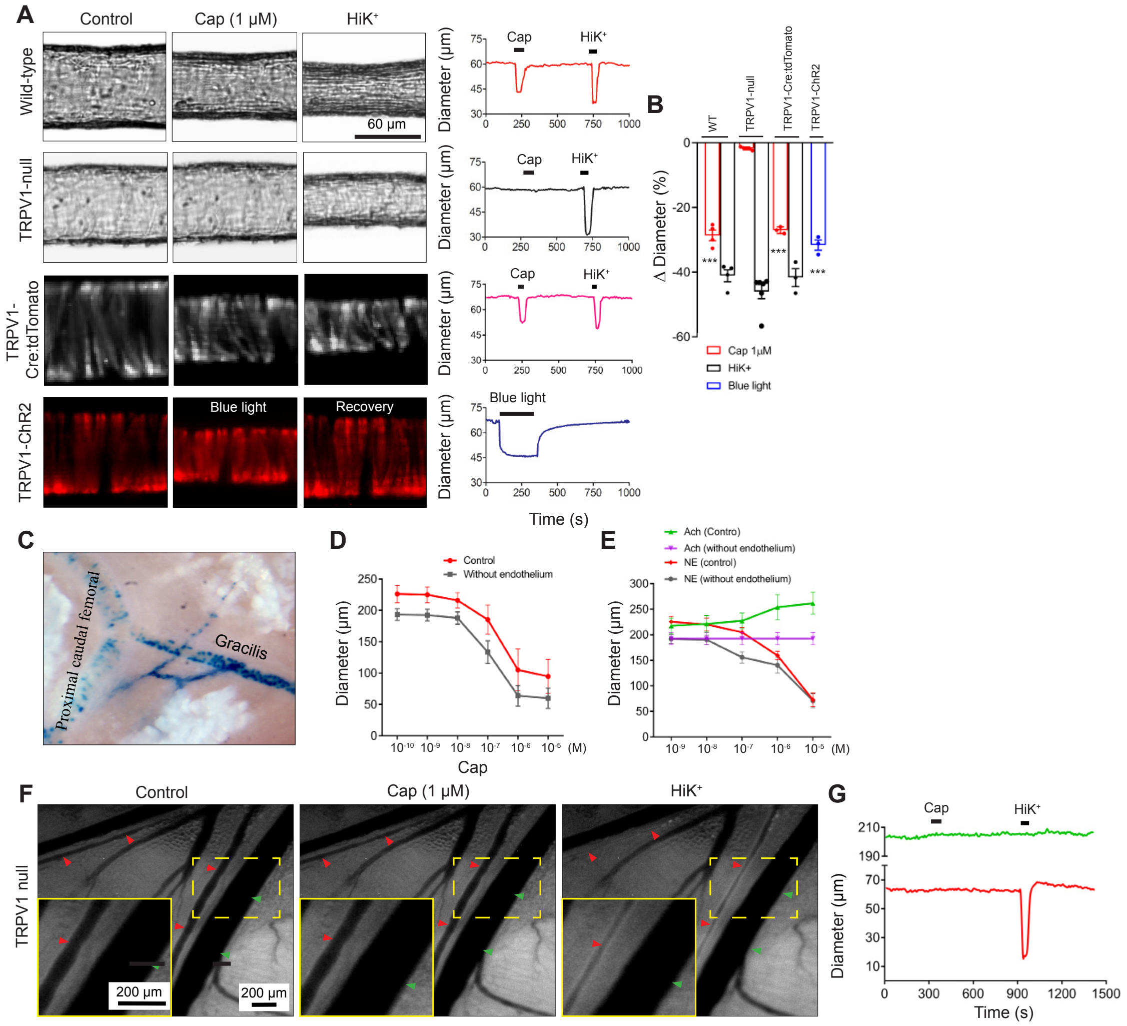


***Figure S4.*** **Arterial contractility in response to TRPV1 agonist and optogenetic stimulation.** (**A**, **B**), Responses of pressurized cerebellar branch arteries from WT, TRPV1-null, TRPV1-Cre:tdTomato and TRPV1-Cre:ChR2 to capsaicin and blue light (n = 4 - 7 arteries from 3 - 5 mice per group, $***P < 0.001$). (**C**)**,** nLacZ staining in gracilis artery of a TRPV1$^{PLAP\text{-}nlacZ}$ mouse. (**D**), Capsaicin-evoked constriction in isolated gracilis arteries with or without endothelium. (**E**), Acetylcholine (ACh) but not norepinephrine (NE) sensitivity is lost in denuded arteries. (**F**, **H**)**,** Intravital imaging of a radial branch artery in a TRPV1-null mice in response to local application of capsaicin and KCl.

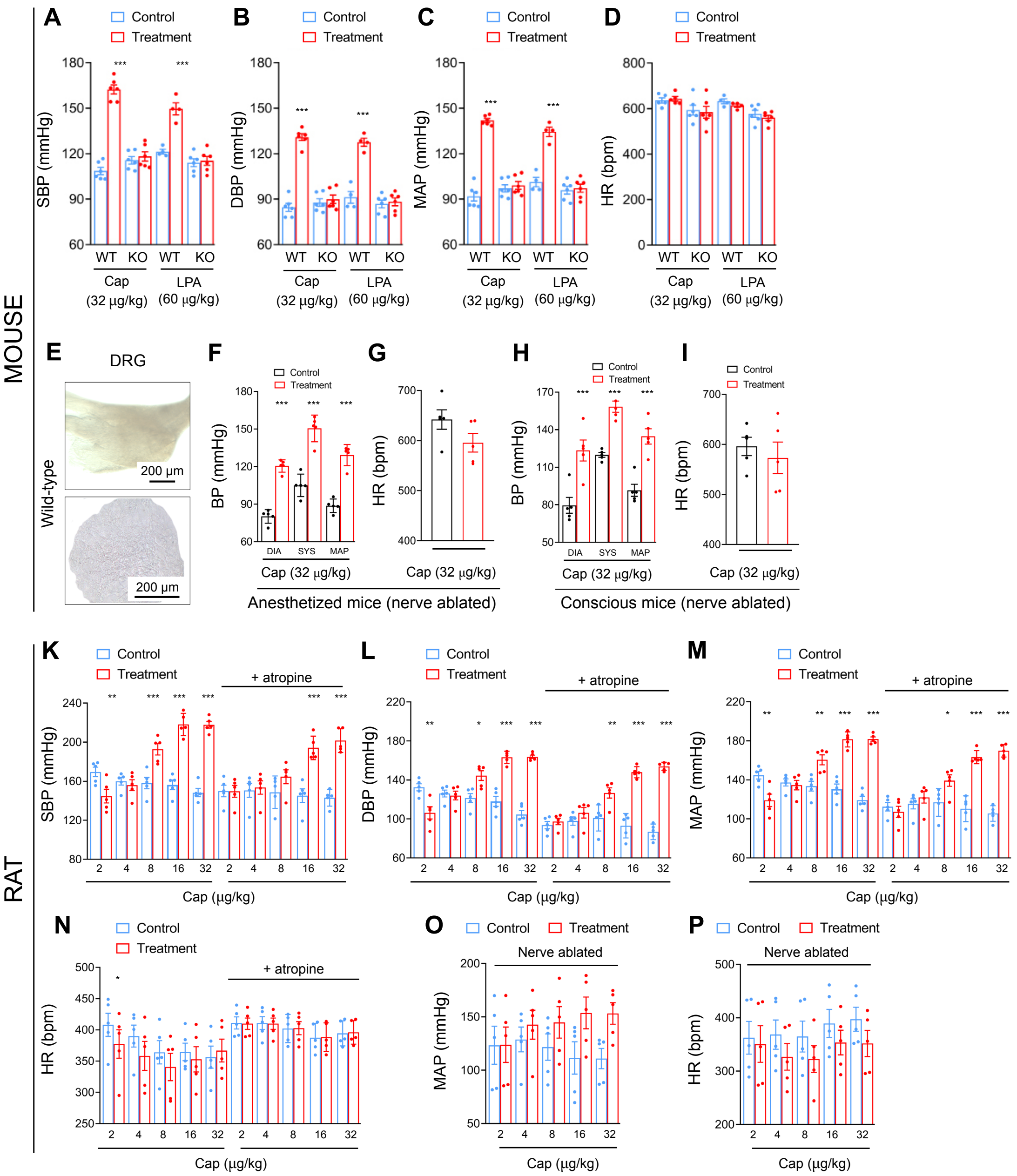


***Figure S5*. Effects of TRPV1 agonists and antagonists on systemic blood pressure**. (**A**-**D**), Systolic BP, diastolic BP, mean arterial pressure and heart rate (HR) values in mice before, and the peak pressor response after, capsaicin administration. (**E**), X-gal staining in control DRG ganglia and slice. (**F**-**I**), BP and heart rate in anesthetized and conscious mice treated with RTX as neonates. (**K**-**N**) BP and heart rate responses (during peak pressor phase) to capsaicin in rats, [Note: capsaicin evokes similar responses with or without atropine pretreatment] (n = 6, **P < 0.05*, *** P < 0.01*, ****P < 0.001*). (**O**, **P**), MAP and HR responses to capsaicin in sensory nerve-ablated rats (neonatal capsaicin treatment, n=5).

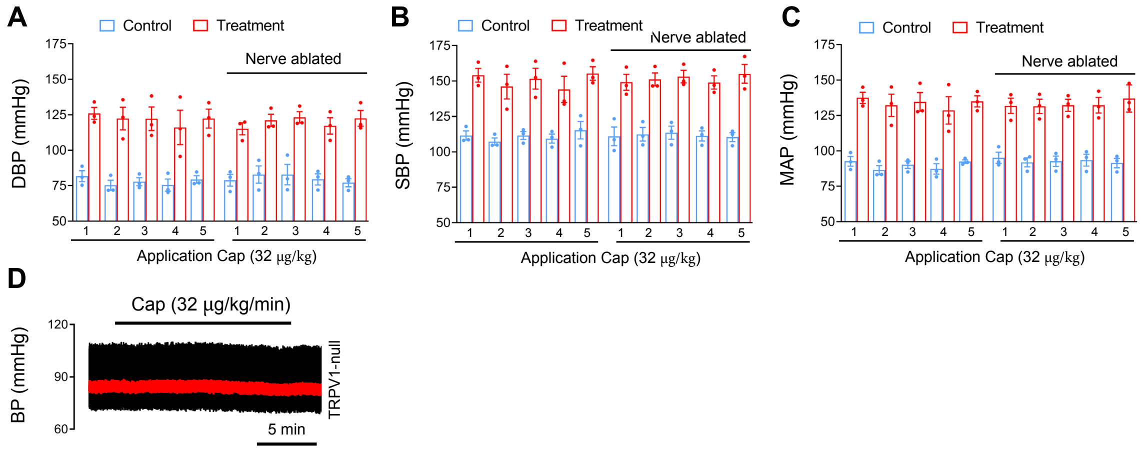


***Figure S6.*** **BP responses to repeated capsaicin administration.** (**A-C**), Diastolic (DBP), systolic (SBP) and mean arterial pressure (MAP) in WT and sensory nerve ablated mice during repeated administration of capsaicin (n=3). (**D**), Capsaicin infusion in TRPV1-null mouse does not alter BP.
